# Chromosome-level genome assembly of the disco clam, *Ctenoides ales*, a first for the bivalve order Limida

**DOI:** 10.1101/2024.03.01.583045

**Authors:** Kyle E McElroy, Rick Masonbrink, Sivanandan Chudalayandi, Andrew J Severin, Jeanne M Serb

## Abstract

The bivalve subclass Pteriomorphia, which includes the economically important scallops, oysters, mussels, and ark clams, exhibits extreme ecological, morphological, and behavioral diversity. Among this diversity are five morphologically distinct eye types, making Pteriomorphia an excellent setting to explore the molecular basis for the evolution of novel traits. Of pteriomorphian bivalves, Limida is the only order lacking genomic resources, greatly limiting the potential phylogenomic analyses related to eyes and phototransduction. Here, we present the first limid genome assembly, the disco clam, *Ctenoides ales*, which is characterized by invaginated eyes, exceptionally long tentacles, and a flashing light display. This genome assembly was constructed with PacBio long reads and Dovetail Omni-C^TM^ proximity-ligation sequencing. The final assembly is ∼2.3Gb and over 99% of the total length is contained in 18 pseudomolecule scaffolds. We annotated 41,064 protein coding genes and report a BUSCO completeness of 91.9% for metazoa_obd10. Additionally, we report a completely annotated mitochondrial genome, also a first for Limida. The ∼20Kb mitogenome has 12 protein coding genes, 22 tRNAs, 2 rRNA genes, and a 1,589 bp duplicated sequence containing the origin of replication. The *C. ales* nuclear genome size is substantially larger than other pteriomorphian genomes, mainly accounted for by transposable element sequences. We inventoried the genome for opsins, the signaling proteins that initiate phototransduction, and found that, unlike its closest eyed-relatives, the scallops, *C. ales* lacks duplication of the rhabdomeric G_q_-protein coupled opsin that is typically used for invertebrate vision. In fact, *C. ales* has uncharacteristically few opsins relative to the other pteriomorphian families, all of which have unique expansions of xenopsins, a recently discovered opsin subfamily. This chromosome-level assembly, along with the mitogenome, will be valuable resources for comparative genomics and phylogenetics in bivalves and particularly for the understudied but charismatic limids.

**Significance:** This high-quality chromosome-level genome assembly for *Ctenoides ales*, the disco clam, is the first genome sequenced from the bivalve order Limida, the only group of Pteriomorphia – a highly diverse and ecologically important clade – lacking genomic resources. The sequence and annotation of the *C. ales* genome will be a useful resource for molluscan phylogenetics and comparative genomics.

## INTRODUCTION

The bivalve subclass, Pteriomorphia, includes many of the most economically important bivalves such as mussels, oysters, and scallops. Among the numerous morphological innovations in this clade are nonhomologous pallial eye types, including: pigmented cups and compound eyes in Arcidae, cap eyespots in Ostreidae, mirror eyes in Pectinidae, and invaginated eyes in Limidae (Audino et al. 2020). The many origins of eyes in Pteriomorphia make this clade a compelling setting to study how novel traits arise and whether divergence in genetic architecture underlies this evolution.

To date, over 30 pteriomorphian genomes have been sequenced including at least one chromosome-level assembly from Arcida (Bai et al. 2019), Mytilida (Yang et al. 2021), Ostreida (Peng et al. 2020; Takeuchi et al. 2022), and Pectinida (Kenny et al. 2020). Outside of scallops (Pectinidae), no other eyed species have been sequenced, therefore limiting the possibility of comprehensive phylogenomic comparative studies on the evolution of eyes in this taxonomic lineage. Here, we present an annotated chromosome-level genome assembly of *Ctenoides ales* (Finlay, 1926), an eyed limid species. This genomic data will be valuable to exploring the repeated evolution of eyes in Pteriomorphia and represents the first genomic data from Limida, a diverse order of about 200 extant species (MolluscBase) commonly referred to as flame scallops, file clams, or file shells.

The charismatic *C. ales* (Figure 1a) is known as the “disco clam” or “electric flame scallop” for its flashing mantle display. Despite its bioluminescent-like appearance, this presumed anti-predator display is actually the result of light reflecting from silica nanospheres incorporated into the mantle tissue (Dougherty et al. 2014). Limids are also known for their brightly colored mantle tissue lining the two valves, which is a source of chemical deterrent from predators (Dougherty et al. 2019). Long tentacles are used not only for sensory perception, but also for swimming and chemical defense. (Mikkelsen and Bieler 2003; Donovan et al. 2004; Dougherty et al. 2019). At the base of these tentacles in many limid species are multiple “invaginated” eyes embedded in the mantle tissue (Bell and Mpitsos 1968; Mpitosos 1973; Morton 2000). Similar to scallops (Gorman and McReynolds 1969), limid eyes have two distinct retinas, a “proximal” and “distal” that are made up of rhabdomeric and ciliary photoreceptor cells, respectively (Speiser et al. 2023), with opposing responses to light (Mpitosos 1973). However, the morphological characterization of limid eyes is complicated by differing interpretations and taxonomically narrow studies. Whether limids have spatial vision is still unknown (Speiser et al. 2023). Morphological and behavioral analyses suggests that *C. ales* has poor visual resolution and is unable to distinguish light directionality (Dougherty et al. 2017), in contrast to their closest eyed relatives, the scallops, equipped with image-forming eyes (Land 1965). Genomic comparisons across a variety of eyed and eyeless pteriomorphians should yield great insight into the molecular evolutionary process underlying the emergence of novel photoreceptive organs.

**Figure 1:**
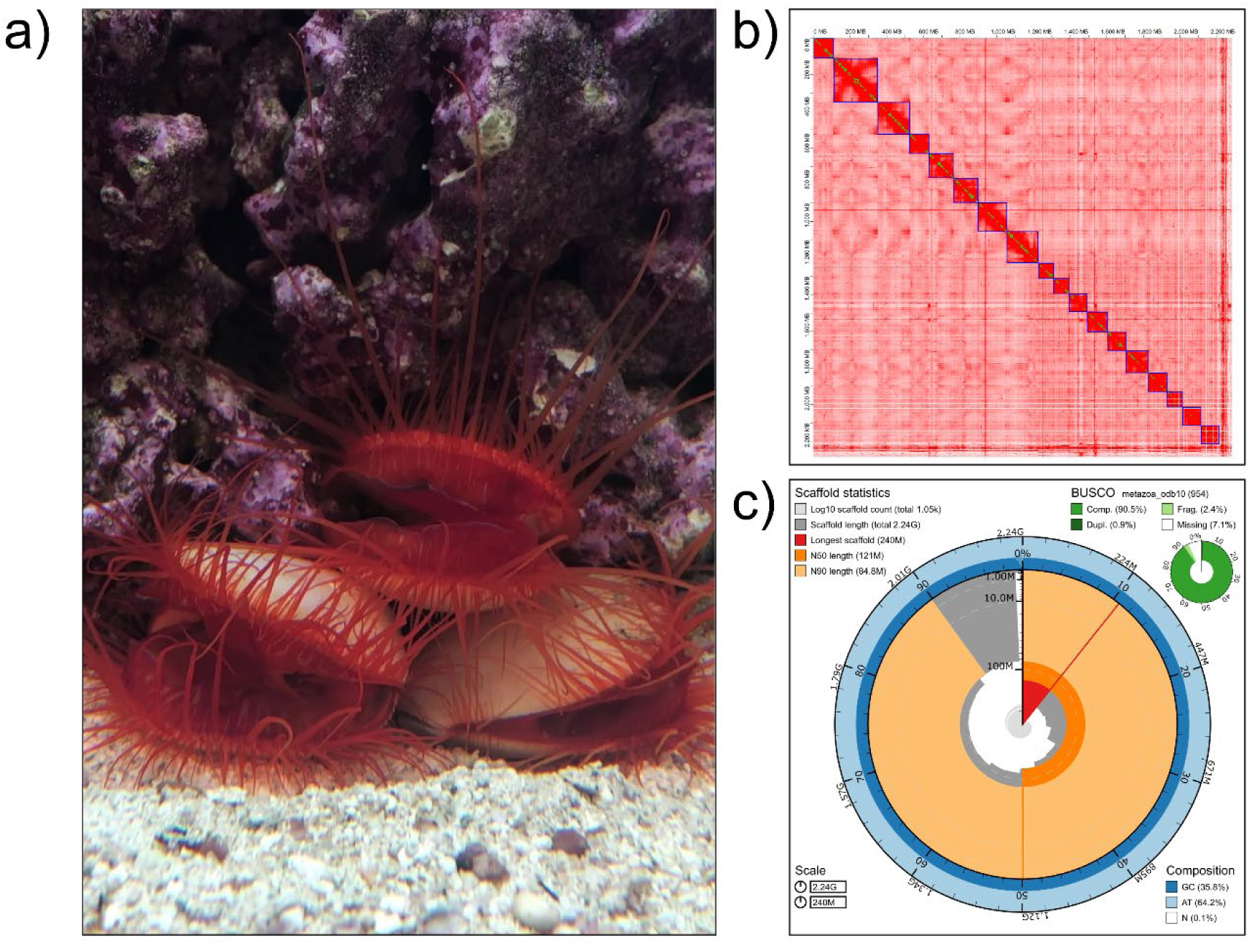
*Ctenoides ales* and summary of its genome assembly. **A)** Adult *Ctenoides ales* in aquarium setting, flashing light display visible in middle individual. Image credit: Jeanne M Serb. **B)** Hi-C contact map for *C. ales*, highlighting the 18 chromosomes recovered from the genome assembly. Darker red indicates higher density of contact, blue and green boxes denote chromosome and contigs, respectively. **C)** Snail plot summarizing key assembly statistics for final *C. ales* assembly with BUSCO results.

Gene duplication is an important source of new genetic information that can be used in the evolution of novel traits (Ohno 1970; Lynch and Conery 2000; Zhang 2003; Birchler and Yang 2022). Opsin duplication is an important for the path to wider spectral sensitivity, particularly in vertebrates (e.g., Escobar-Camacho et al. 2020; reviewed in Hagen et al. 2023) and arthropods (Briscoe 2001; Briscoe et al. 2010; Bentley et al. 2016). Opsins are transmembrane G-protein coupled receptor (GPCR) proteins that form photopigments by binding to a chromophore, typically a vitamin A-derived retinol, which upon light-absorption starts a phototransduction cascade. Opsins are a phylogenetically diverse and widespread protein family with a variety of both ocular and extraocular light-dependent functions, and light-independent functions (Terakita 2005; Shichida and Matsuyama 2009; Terakita and Nagata 2014; Moraes et al. 2021). Opsin classification is based on the specific G-protein an opsin activates (*e.g.*, G_q_, G_t_, G_i_, G_o_, G_s_), the type of photoreceptor cell where it is expressed (*e.g.*, ciliary vs. rhabdomeric), and its phylogenetic placement (*e.g.*, tetraopsins) (reviewed in Shichida and Matsuyama 2009; Porter et al. 2011). Though the phrase “visual opsin” has recently come into question (Feuda et al. 2022), typically vertebrates rely on ciliary (c-type) opsins that couple with G_t_ proteins for vision, bilaterian invertebrates use rhabdomeric (r-type) G_q_-opsins, and cnidarians use a distinct group of G_s_-opsins. Transcriptome sequencing has revealed multiple duplications of the rhabdomeric opsins (r-opsins) in scallops (Porath-Krause et al. 2016), which have a complex, image-forming mirror eye, raising the possibility that opsin duplication is characteristic of eyed pteriomorphian lineages. We explore this hypothesis by scanning the *C. ales* genome assembly for opsins and phylogenetically analyzing them in the context of opsin evolution in Pteriomorphia.

## MATERIALS AND METHODS

### Sample collection and DNA extraction

Live *C. ales* individuals were dissected from their shells, immediately flash frozen in liquid nitrogen and stored at -80C. Frozen adductor muscle tissue was homogenized into a fine power and DNA was then extracted via a cetyltrimethylammonium bromide (CTAB) protocol. Briefly, powdered tissue was suspended in a β-mercaptoethanol and CTAB buffer solution, then incubated at 68°C with occasional agitation followed by incubation at 60°C with RNAse A and Proteinase K. After centrifugation, the aqueous layer underwent phenol/chloroform extraction twice, then a single chloroform/isoamyl alcohol extraction. DNA was then precipitated with 0.7 volumes of isopropanol. The precipitated DNA was resuspended in Qiagen G2 buffer with RNase A and Qiagen Protease and incubated at 50°C with occasional agitation, and finally, vortexed and centrifuged to pellet debris.

### PacBio library and sequencing

DNA samples were quantified using Qubit 2.0 Fluorometer (Life Technologies, Carlsbad, CA, USA). The PacBio SMRTbell library (∼20kb) for PacBio Sequel was constructed using SMRTbell Express Template Prep Kit 2.0 (PacBio, Menlo Park, CA, USA) using the manufacturer’s recommended protocol. The library was bound to polymerase using the Sequel II Binding Kit 2.0 (PacBio). Sequencing was performed on PacBio Sequel II 8M SMRT cells.

### Dovetail Omni-C library preparation and sequencing

For the Dovetail Omni-C library, chromatin was fixed in place with formaldehyde in the nucleus and then extracted. Fixed chromatin was digested with DNAse I, chromatin ends were repaired and ligated to a biotinylated bridge adapter followed by proximity ligation of adapter containing ends. After proximity ligation, crosslinks were reversed, and the DNA purified. Purified DNA was treated to remove biotin that was not internal to ligated fragments. Sequencing libraries were generated using NEBNext Ultra enzymes and Illumina-compatible adapters. Biotin-containing fragments were isolated using streptavidin beads before PCR enrichment of each library. The library was sequenced on an Illumina HiSeqX platform to produce approximately 30x sequence coverage. Then HiRise used MQ>50 reads for scaffolding.

### RNA isolation and sequencing

Flash frozen whole-animal tissue was stored at -80C until preparation for RNA isolation. For every animal used, a sample of adductor, mantle, and eyes was prepared. Isolated tissue was first ground into a homogenous powder using an OPS Diagnostics LLC CryoGrinder^TM^ System and then stored at -80°C until RNA isolation. Total RNA was isolated from ground tissue powder using an E.Z.N.A Total RNA Kit II (Omega BIO-TEK). RNA purity was assessed with Nanodrop. RNA integrity and quantity was determined with an Agilent 2100 Bioanalyzer (Agilent Technologies). Libraries for sequencing were prepared with the NEBNext Ultra Directional RNA Library Prep Kit (Illumina), following poly(A) mRNA enrichment with Oligo dT Beads, and then sequenced (2×150 bp reads) on an Illumina NovaSeq 6000 at Iowa State University’s DNA Facilty.

### Draft assembly

Approximately 290.8 Gb of PacBio CLR reads were used as an input for assembly by WTDBG2 v2.5 (Ruan and Li 2020) with genome size set to 1.7 Gb, minimum read length 20,000, and minimum alignment length 8,192. Additionally, realignment was enabled with the -R option and read type was set with the option -x sq. Blast results of the WTDBG2 output assembly against the nt database were used as input for blobtools v1.1.1 (Laetsch and Blaxter 2017) and scaffolds identified as possible contamination were removed from the assembly. Finally, purge_dups v1.2.3 (Guan et al. 2020) was used to remove haplotigs and contig overlaps.

### HiRise scaffolding

The de novo WTDBG2 assembly and Dovetail Omni-C library reads were used as input data for the proximity ligation-based genome-scaffolding pipeline, HiRise (Putnam et al. 2016). Dovetail Omni-C library sequences were aligned to the draft input assembly using BWA (https://github.com/lh3/bwa). The separations of Dovetail Omni-C read pairs mapped within draft scaffolds were analyzed by HiRise to produce a likelihood model for genomic distance between read pairs, and the model was used to identify and break putative misjoins, to score prospective joins, and make joins above a threshold.

### Final assembly

Dovetail assembly scaffolds were scaffolded using Juicer v1.5.7 (Durand et al. 2016), 3D-DNA v180114 (Dudchenko et al. 2017), Juicebox v1.11.08 (Robinson et al. 2018), with a BWA 0.7.17 alignment. Blobtools v2.2.0 was used to identify putative contaminants using BLAST+ v2.11.0 (Camacho et al. 2009) to NCBI nt database (downloaded May 30, 2020) and alignments of PacBio subreads with Minimap v2.2 (Li 2018). Contigs that could not be placed in pseudomolecules were subjected to redundancy filtering using coordinates from two criteria: mapping contigs to the pseudomolecules with Minimap v2.2 and repeats identified as described below. These coordinates were merged using Bedtools (Quinlan and Hall 2010) merge, and when a contig could achieve 90% identity across 90% of the contig length to a pseudomolecule it was removed. Genome completeness was assessed using BUSCO v5.1.2 (Waterhouse et al. 2018) to Metazoa_odb10 and Eukaryota_odb10.

### Transposable element characterization

De novo repeat identification was conducted with RepeatModeler v2.0.2 (Flynn et al. 2020), utilizing RepeatScout v1.0.6 (Price et al. 2005), RECON v1.08 (Bao and Eddy 2002), and LTR_Retriever v2.9.0 (Ou and Jiang 2018) and the Extensive de-novo TE Annotator (EDTA) v2.0.1 (Ou et al. 2019), which uses three pipelines for TE discovery based on structural characteristics of terminal inverted repeated (TIR) DNA transposons, long terminal repeat (LTR) retrotransposon, and helitrons. TEsorter v1.4.6 (Zhang et al. 2022) was used next to classify sequences from the RepeatModeler and EDTA output based on REXdb v. 3 HMM profiles and combined classified sequences and reduced redundancy between the two sets of repeats with CD-HIT-EST v4.8.1 (Li and Godzik 2006; Fu et al. 2012). Because SINEs lack any coding sequences and no structural detection pipeline is included in EDTA, the sequences annotated as SINEs by RepeatModeler were queried against Repbase online (Bao et al. 2015) with CENSOR and kept sequences matching SINEs with scores >250. We combined the TEsorter classified sequences with SINEs and the remaining EDTA sequences identified via structural features (e.g., LTRs) into a single TE library for each species masked each genome with its specific TE library using RepeatMasker v4.1.2-p (Smit et al.). We used the RepeatMasker script buildSummary.pl to summarize the TE content of each genome and calcDivergenceFromAlign.pl to collect the Kimura substitution levels of TE copies to generate a repeat landscape. For comparison, we performed these analyses on the *C. ales* genome and other available pteriomorphian genome assemblies (see below).

### Gene prediction and functional annotation

RNA-seq reads were aligned to a softmasked genome using Star v2.5.3a (Dobin et al. 2013). BUSCO v5.1.2 was used to train Augustus v3.3.2 (Stanke and Waack 2003; Stanke et al. 2006) with Metazoa_odb10 to use for gene prediction with Braker v2.1.2 (Brůna et al. 2021) and GeneMark v4.38 (Brůna et al. 2020). Genomethreader v1.7.3 (Gremme et al. 2005) was used to align 283,363 proteins downloaded from Uniprot and delimited by the Bivalvia taxon. Using both datasets, the gene annotations were further refined using Mikado v2.3.2 (Venturini et al. 2018) with Transdecoder v5.5.0 (Haas, https://github.com/TransDecoder), Portcullis v1.2.2 (Mapleson et al. 2018) and BLAST+ v2.11.0. Functional gene annotations were created via Diamond v2.0.4 (Buchfink et al. 2015) searches to NCBI NR (downloaded May 3, 2021), Uniprot/Swissprot (downloaded May 28, 2022), and Interproscan v5.38 (Jones et al. 2014).

### Phylogenetic analysis

Protein sequences were collected from annotated bivalve genomes, including 11 other pteriomorphian species and two outgroups (*Mercenaria mercenaria* and *Sinonovacula constricta*) to phylogenetically compare the protein coding-content of the *Ctenoides ales* genome (Table S1). A total of 508,333 proteins were analyzed with OrthoFinder v2.5.4 (Emms and Kelly 2019). From these results, 1,156 proteins were identified as single-copy and present in all 14 species analyzed. We used these protein sequences to produce a species-tree. The amino acid sequences identified as single copy and complete for all 14 species analyzed using mafft v7.481 (--auto)(Katoh et al. 2002; Katoh and Standley 2013). Next, we trimmed the alignments with trimal v1.4.rev15 (-automated1) (Capella-Gutiérrez et al. 2009). Then we used IQtree2 v2.1.3 (Nguyen et al. 2015; Minh et al. 2020) with modelfinder (Kalyaanamoorthy et al. 2017) to generate maximum likelihood trees for each trimmed protein alignment and a summary file of protein substitution models used. The substitution model results were combined with a gene partition file generated with catsequences (V1.3; 10.5281/zenodo.4409153) and used as input for a partitioned IQtree2 ML analysis (Chernomor et al. 2016). Branch support was evaluated by ultrafast bootstrap (Hoang et al. 2018), SH approximate likelihood ratio test, and approximate Bayes test features in IQtree2 (-B 1000 --alrt 1000 --abayes) (Anisimova et al. 2011).

A second species tree was generated to include multiple taxa per pteriomorphian family where high-quality genomes were available but not genome annotations (i.e., additional Arcidae). This was done to account for variation within taxonomic families in downstream characterization of transposable element and opsin content. As in McElroy et al. (2023), BUSCO was used to predict conserved protein sequences to construct a species tree independent of genome annotations. We downloaded genomes assemblies from 12 pteriomorphian species, three representatives from the families Arcidae, Pectinidae, Ostreidae, and Mytilidae, along with four other bivalve species as outgroups (accessions and assembly statistics listed in Table S2). We ran BUSCO v5.2.2 on each of these 16 genome assemblies using the metazoa_odb10 database (BUSCO scores listed in Figure S1, Table S2). We then used the 177 BUSCO amino acid sequences identified as single copy and complete for all 16 species as input for a maximum-likelihood analysis following the same methods described for the first species tree.

### Synteny between *C. ales* and *P. maximus*

Synteny was determined using i-ADHoRe v3.0.01 (Proost et al. 2012) using the OrthoFinder2 results and their respective assemblies and annotations. Synteny was only performed for scaffolds larger than 1 Mb. The following parameters were included in the i-ADHoRe config: blast_table=black.blastTable, prob_cutoff=0.001, anchor_points=3, number_of_threads=36, visualizeAlignment=false, output_path=out_5, alignment_method=nw, gap_size=25, cluster_gap=50, level_2_only=true and q_value=0.9. The function dashbio.Circos in the dashbio library was used to visualize the synteny in a circos plot.

### Opsin identification and analysis

We identified opsins via Phylogenetically Informed Annotation tool (PIA; Speiser et al. 2014) from the *C. ales* gene models and the homology-based *de novo* gene prediction with the BITACORA pipeline v1.3 (Vizueta et al. 2020) using GeMoMa (Keilwagen et al. 2016; Keilwagen et al. 2018) and a database of molluscan opsin sequences described in McElroy et al. (2023). For PIA, we used the modified version from https://github.com/MartinGuehmann/PIA2 and the Light Interacting Toolkit (LIT_1.1; opsins using r_opsin_20_rtrans.fas for opsin classification). For further opsins classification and phylogenetic comparison, we included McElroy et al. (2023) opsins for the following species: *Argopecten irradians*, *Pecten maximus*, *Mizuhopecten* (*Patinopecten*) *yessoensis*, *Scapharca broughtinii*, *Scapharca kagoshimensis*, *Tegillarca granosa*, *Perna viridis*, *Mytilus coruscus*, *Mytilus galloprovincialis*, *Crassostrea gigas*, and *Crassostrea virginica*. We additionally generated opsin models for *Ostrea edulis*. For all opsins, we manually inspected the sequences to ensure high quality (i.e., intact GPCR Class A 7tm_1 domain and containing the K296 position). For outgroup sequences, we used melatonin receptors and the opsin-like sequences from Placozoa, “placopsins” (Feuda et al. 2012), along with the more recently described GPRC relative of opsins found in lophotrochozoans, the “pseudopsins” (De Vivo et al. 2023). We aligned the opsin and outgroup amino acid sequence for the 13 pteriomorphian species using mafft [--maxiterate 1000 --genafpair]. We then generated a maximum likelihood tree in IQtree2 with modelfinder (best-fit model Q.yeast+F+R8 according to Bayesian Information Criterion) that was evaluated with ultrafast bootstrap, SH approximate likelihood ratio test, and approximate Bayes test features in IQtree2 (-B 1000 --alrt 1000 --abayes).

### Mitogenome assembly and assembly

To characterize the mitochondrial genome, a partial *C. ales* cytochrome c oxidase subunit I (COI) (MF540379.1) was queried against the final genome assembly using blastn. This search yielded no returns. Next, the draft, unfiltered WTDBG2 assembly was searched, returning a single contig approximately 18 kb long containing the partial COI sequence. The PacBio reads were mapped to the mt contig with minimap2 v2.14-r883 (Li 2018). Aligned reads were then extracted and assembled with flye v2.9 (Kolmogorov et al. 2019), which generated a single circular contig 20,859 bp long. This contig was then annotated on the MITOS (Genetic Code 5: Invertebrate Mitochondrial) (Bernt et al. 2013) and MITOS2 (Donath et al. 2019) web servers (RefSeq 89 Metazoa; Genetic Code 5: Invertebrate Mitochondrial). Additionally, the mitochondria genome was scanned for open reading frames (ORFS) with OFRfinder on NCBI (https://www.ncbi.nlm.nih.gov/orffinder/) (Genetic Code 5: Invertebrate Mitochondrial; “ATG” and alternative start codons). The ORFfinder results were used to characterize the complete coding sequences for each of the 12 proteins identified. ARWEN v1.2 (Laslett and Canbäck 2008) was additionally used to evaluate tRNAs predicted by MITOS and potentially identify the two tRNAs classified as “missing” by MITOS annotation. The PacBio and forward reads from the Omni-C Illumina sequences were mapped (minimap and bwa-mem2, respectively) to evaluate the apparent duplication of the OH sequence. For the Illumina mapping, a version of the mitogenome with only one copy of the duplicated region was also used as a reference to compare coverage over that sequence.

## RESULTS AND DISCUSSION

### Genome assembly and completeness analysis

Dovetail performed a de novo genome assembly of *Ctenoides ales*. Approximately 290 billion bases of PacBio Continuous Long Reads (CLR) were used in the assembly (14,188,342 reads, average length = 20.5Kb). The WTDBG2 assembler generated a draft assembly, which was subsequently scaffolded with 602 million Omni-C reads (Table S3 for intermediate stats). These reads were then used to further scaffold the Dovetail genome using Juicer, 3D-DNA, and manually corrected using Juicebox. After eliminating haplotigs and contaminants, the final assembly contained 1,049 scaffolds and 2.2 billion bases. Among these scaffolds, the 18 largest correspond to chromosomes, accounting for 99.16% of the total nucleotide content (Figure 1b). The assembly’s scaffold N50 value is 121 Mb, with the longest scaffold measuring 240 Mb (Figure 1c). The completeness of this assembly is exemplified by the 98.56% mapping rate of the PacBio long reads against it, which amounts to approximately 100x coverage. Furthermore, the assembly exhibits a BUSCO completeness score of 91.9% using the metazoa_odb10 gene set (Figure 1c) and an 85.4% score based on the mollusca_odb10 gene set. This BUSCO-based assembly quality is in line with other pteriomorphian assemblies (Figure S1), including scallops, however the genome is considerably larger. For example, the *Pecten maximus* genome released in 2019 by the Wellcome Sanger Institute is chromosomal with 19 chromosomes and 3,983 scaffolds containing 918 million bases (Kenny et al. 2020), making our *C. ales* genome assembly over double in size (see Table S2 for assembly size comparisons). We further examined the genome size disparity between *C. ales* and other pteriomorphians in the context of gene content, genome duplication, and transposable elements.

### Gene content and phylogenetic analysis

RNA-seq data were generated at the Iowa State University DNA Facility using eye, mantle, and adductor muscle tissues from 10 *C. ales* specimens. Gene prediction yielded 41,064 gene models with an average gene length of 23,950 bp (Table 1 for further details). We aligned amino acid sequences from predicted genes against the NR and Swissprot databases and assigned functional information to 77.7% of genes. The metazoan BUSCO scores of the gene annotation indicate a high degree of completeness at 89.2%, which is qualitatively similar to the results from the genome assembly. These gene model predictions suggest that *C. ales* may have nearly 50% more genes than is typical for some pteriomorphians. The first scallop genomes, *Mizuhopecten* (*Patinopecten*) *yessoensis* and *Chlamys farreri* were described as having 26,415 (Wang et al. 2017) and 28,602 (Li et al. 2017), respectively. While the scallop *Pecten maximus* was initially reported to have 67,741 protein coding genes (Kenny et al. 2020), Zeng et al. (2021) identified 26,995 genes. Arcidae gene reports tend to have similar numbers of genes as scallops, e.g., 24,045 in *Scapharca broughtonii* (Bai et al. 2019) and 24,398 in *Tegillarca granosa* (Bao et al. 2021). The number of protein coding genes predicted in *C. ales* is more comparable to genome annotations from Mytilidae and Ostreidae genomes, which tend to contain between 30,000-40,000 genes (e.g., Peñaloza et al. 2021; Yang et al. 2021). Notably, transposable elements (TEs) do not appear to make up a substantial portion of the predicted genes as only 1,825 genes (∼4% of total) have annotation terms from likely TEs (“transposase”, “reverse transcriptase”, “helicase”, “LINE”, “integrase”, “RNAse H”). Therefore, the relatively high number of genes predicted in this genome are not likely inflated by TEs annotated as genes. Variation in gene model prediction strategies may account for some of the differences in gene totals, as fragmentation of lowly expressed genes may have occurred. Additionally, factors such as genome size and evolutionary processes may contribute to these differences.

**Table 1:** Summary statistics of *Ctenoides ales* genome & annotation.

| **Assembly stats** |  | **Annotation stats** |  |
| --- | --- | --- | --- |
| Number of scaffolds | 1,049 | Number of genes | 41,064 |
| Length of assembly | 2,237,305,496 bp | Number of mRNA | 44,726 |
| Longest scaffold | 240,195,547 bp | Mean gene length | 23,950 bp |
| Scaffold L50/N50 | 7/121,434,358 bp | Mean CDS length | 1,346 bp |
| Scaffold L90/N90 | 16/84,817,633 bp | Mean exon per gene | 7.2 |
| Scaffold CG Content | 35.77% | Mean introns per gene | 6.2 |
| Scaffold N content | 0.09% | Mean exon length | 188 bp |
| Number of contigs | 8,191 | Mean intron length | 4,375 bp |
| Longest contig | 7,630,288 bp | Total CDS length | 50,641,044 bp |
| **Contig L50/N50** | 508/1,207,478 bp |  |  |
| **Contig L90/N90** | 2476/ 133,914 bp |  |  |

**Table 2:**
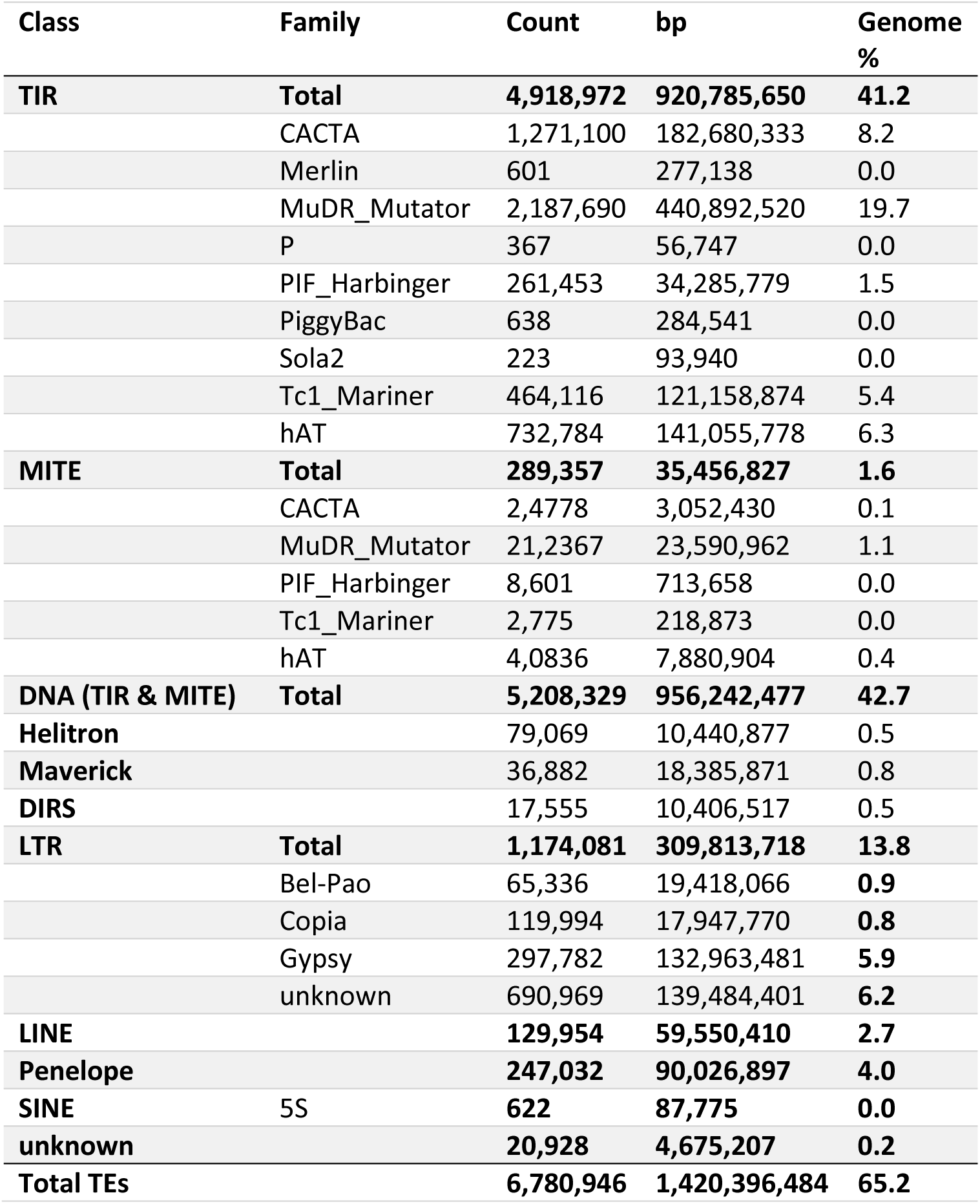
Transposable element content in *Ctenoides ales* genome.

To phylogenetically compare the protein coding-content of the *Ctenoides ales* genome, we collected protein sequences from annotated bivalve genomes, including 11 pteriomorphians and two outgroup species (*Mercenaria mercenaria* and *Sinonovacula constricta*). We analyzed a total of 508,333 proteins with OrthoFinder2 (Emms and Kelly 2015; Emms and Kelly 2019), which resulted in clustering 91.5% of proteins into 465,163 orthogroups (summarized in Table S4). Only 7.4% of the proteins belong to species-specific orthogroups. For *C. ales*, 33,033 out of the 39,079 protein sequences analyzed were placed into orthogroups, 5,230 of which were specific orthogroups to *C. ales*, meaning that the majority (71.1%) of protein-coding genes from our annotation are shared with other species (e.g., Figure S2). A lack of close relatives may have influenced these results, as species with the highest proportion of genes in species-specific orthogroups are those that are the sole representative of a family (*C. ales*: 13.4%, *M. mercenaria*: 19.5%, *P. fuctata*: 16.3%) We used the 1,156 single-copy orthologs found in all 14 species to reconstruct the species phylogeny using maximum-likelihood analysis with IQ-TREE2. This phylogeny had high support values at all nodes and met expectations of species relationships, including *C. ales* representing Limidae as a sister lineage to Pectinidae (e.g., Audino et al. 2020) (Figure 2b). Together, these analyses of the *C. ales* gene content are indicative of a high-quality annotation for this first limid genome. Furthermore, though the gene count for *C. ales* is relatively high within Pteriomorphia, protein coding sequence accounts for 50Mb, or 2.3%, of the total genome assembly length, therefore contributing very litle to its large genome size (e.g., the scallops *P. maximus* and *Mi. yessoensis* have about 41Mb of coding sequence, each). More genome assemblies from limid species will be needed to clarify if the *Ctenoides* genome is broadly representative of species in Limida for gene content and determine whether gene expansions in this lineage predate, accompanied, or followed its genome size increase.

**Figure 2:**
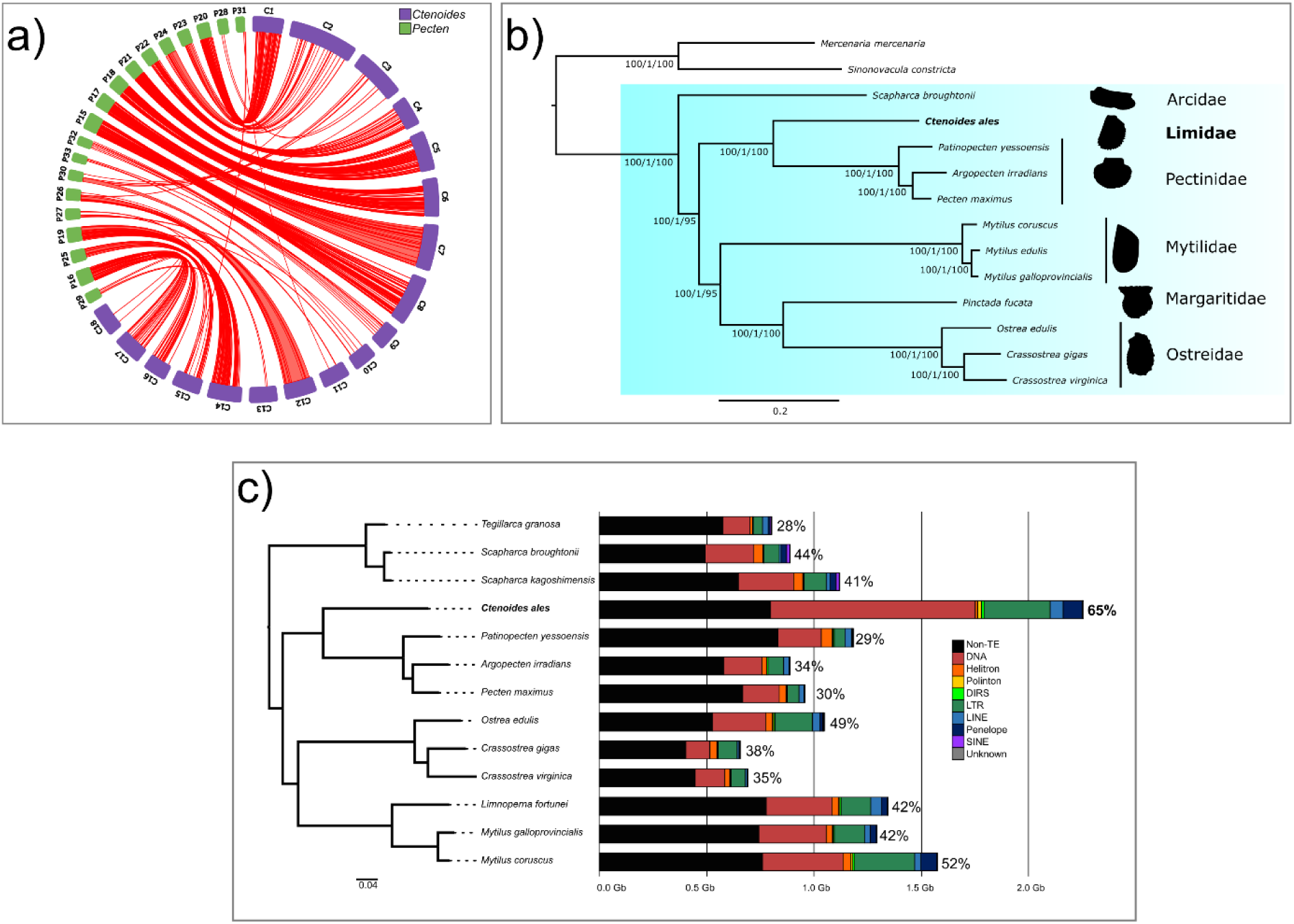
Comparative genomics with other bivalves. **a)** Circos plot of syntenic regions (red arcs) between *C. ales* (in purple) and *P. maximus* (in green). Scaffold names were shortened to their unique numbering and the name portion replaced with a C for *Ctenoides* or a P for *Pecten*. **b)** Phylogenetic placement of *Ctenoides ales* among bivalves. Maximum likelihood species tree generated with IQtree2 based on a partitioned amino acid supermatrix from 1,156 single copy orthologs identified from OrthoFinder2 recovered from all 14 species displayed. Branch values are SH-aLRT % support (with 1,000 replicates)/aBayes probability/UFBoot support % (with 1,000 replicates). **c)**. Genome assemblies for 13 pteriomorphian species characterized by TE amount by distinct types of TEs and “non-TE” portion of genome in black; % given per species is the genomic proportion of TEs.

### Synteny shows no whole genome duplication events

To investigate the potential involvement of whole genome duplication (WGD) in the larger genome size of *Ctenoides ales* than scallops and other pteriomorphian species, we conducted a synteny comparison with the king scallop, *Pecten maximus*. The expectation was to observe a majority of chromosomes exhibiting synteny from one chromosome in *P. maximus* to two chromosomes in *C. ales*, indicating a WGD event. We found no evidence of WGD. Instead, the results revealed a one-to-one correspondence for 12 chromosome pairs (*Ctenoides (C)*:*Pecten (P)*): C2:P24, C3:P30, C4:P22, C5:P21, C6:P18, C7:P17, C8:P15, C9:P32, C12:P26, C14:P19, C15:P25, and C18:P29 (Fig. 2a).

Considering *C. ales* possessing one less chromosome, a fusion event was anticipated. The circos plot demonstrated C1 mapping to P20 and P23, but an inverse pattern was also identified where C15 and C17 both mapped to P16 (Fig. 2a). This observation highlights the occurrence of genomic rearrangement since the divergence of these species from their common ancestor. Genomic rearrangement is further supported by the diminished or fragmented synteny observed between chromosomes C10, C11, and C13 in *C. ales* and P27, P28, P31, and P33 in *P. maximus* (Fig. 2a). The circos plot’s syntenic relationships reveal a high degree of synteny between the two species, accompanied by some genomic rearrangement. However, there was no evidence of a WGD event to account for the substantially larger genomic content in *C. ales*.

In general, high degrees of synteny have been observed across much of Pteriomorphia, including between scallop species (Han et al. 2022), between scallop and ark clam genomes (Bao et al. 2021), and scallop versus Mytilidae comparisons (Yang et al. 2021). The most extensive genome rearrangements in Pteriomorphia are in the oysters, which have notably fewer chromosomes (e.g., Gundappa et al. 2022) than groups such as scallops. Genomic evidence of past WGD has been reported from pteriomorphian genomes (Corrochano-Fraile et al. 2022), but these duplication events have not been placed phylogenetically, making it unclear whether these apparent WGD are shared across Pteriomorphia or occurred in parallel among separate lineages. Our analysis of the first limid genome indicates that no genome duplication has occurred in either Pectinidae or Limidae since their split. Furthermore, while the chromosomes of *C. ales* are much larger than *P. maximus*, gene order has largely been preserved between these lineages.

### Transposable elements contribute to large genome size in *Ctenoides*

The *C. ales* genome is twice as large as genomes from the Pectinidae (Figure 2a), despite no evidence of whole genome duplication. To determine if the size difference was due to repetitive elements, we characterized repeat content from *C. ales* and 12 other pteriomorphians (Table S5) with RepeatModeler, EDTA, and RepeatMasker. We found that much of the difference in genome size across Pteriomorphia is attributable to transposable elements. Based on our estimation, about 65% of the *C. ales* genome is made up of TEs, with DNA transposons making up 40% of the genome. The TE content of *C. ales* is about twice that of the Pectinidae, and greater than any of the 12 other pteriomorphian genomes analyzed here (Figure 2a). The repeat landscape of the *C. ales* genome reveals a large, ancient burst of TE activity, mainly in DNA transposons and more recent burst in LTR *Gypsy* elements (Figure 2b). We also observed far more “intact” (i.e., full structural and coding components) long terminal repeat (LTR) elements from the EDTA analysis than the other species analyzed, which also reflects recent activity of these TEs, such that they have not degraded or been excised from the genome (Table S6).

Several classes of TEs are far more abundant in *C. ales* than other pteriomorphians, particularly DNA transposons and LTR retrotransposons, and, to a lesser degree, LINEs, and Penelope retrotransposons. Helitrons are less abundant in *C. ales* than other species analyzed here (Table S5). SINEs appear to have had very little activity outside of Arcida (Figure 2d, Figure S3), including *C. ales*. Based on the repeat landscapes, there has likely been greater TE activity more recently in the *C. ales* genome than *P. maximus* (Figure S3). We generally observed similar TE content within families, e.g., in Pectinidae *A. irradians*, *P. maximus*, and *Mi. yessoensis* differed by 5% across the three species. Within-family differences in TE content still typically varied less between species in the same genus, e.g., *Crassostrea*, but *M. galloprovincialis* appears to have more TEs than *M. coruscus* (Figure 2a). With *C. ales* being the first limid genome sequenced, it will be important to sequence more broadly from Limida to have a thorough account of how and when this genomic expansion occurred.

TEs are important drivers of genome size evolution (Kidwell 2002) and can account for drastic variation in genome size across closely related species (e.g., Naville et al. 2019; Wong et al. 2019), but the factors influencing differential accumulation and loss of TEs across taxa is still an ongoing area of research. Population genetic theory (Lynch 2007) and empirical data (e.g., Szitenberg et al. 2016) point toward genetic drift as a powerful force influencing TE accumulation. Therefore, changes in effective population size could explain differences in TE content across taxa. Other factors, such as decreased TE silencing (Liu et al. 2022) may also contribute to TE expansion. The genome expansion we found in *C. ales* along with recent evidence for highly variable TE content across bivalves (Martelossi et al. 2023) highlight the importance of this genomically understudied group of animals for exploring the genomic and biological influences on TE evolution.

### Relatively few opsins in the *C. ales* genome

Using the Phylogenetically Informed Analysis tool with the light-detection toolkit (Speiser et al. 2014), we identified 6 opsins from our genome-wide annotation (1 of each: canonical r-opsin, noncanonical r-opsin, xenopsin, G_o_-opsin, retinochrome, and neuropsin) and an additional 2 complete xenopsins from the BITACORA pipeline output. We also found 2 partial opsin sequences (a noncanonical r-opsin and a G_o_-coupled opsin) from the BITACORA output containing the K296 residue but a truncated seven-transmembrane protein domain. To add evolutionary context for these *C. ales* opsins, we generated a ML phylogenetic tree that included opsins from 12 other pteriomorphian species (Figure 3a, Figure S4). As outgroup proteins in this analysis, we included melatonin receptors and the placozoan opsin-like sequences “placopsins” (Feudea et al. 2012), along with the recently described group of closely related GPCRs in lophotrochozoans, “pseudopsins” (De Vivo et al. 2023).

**Figure 3:**
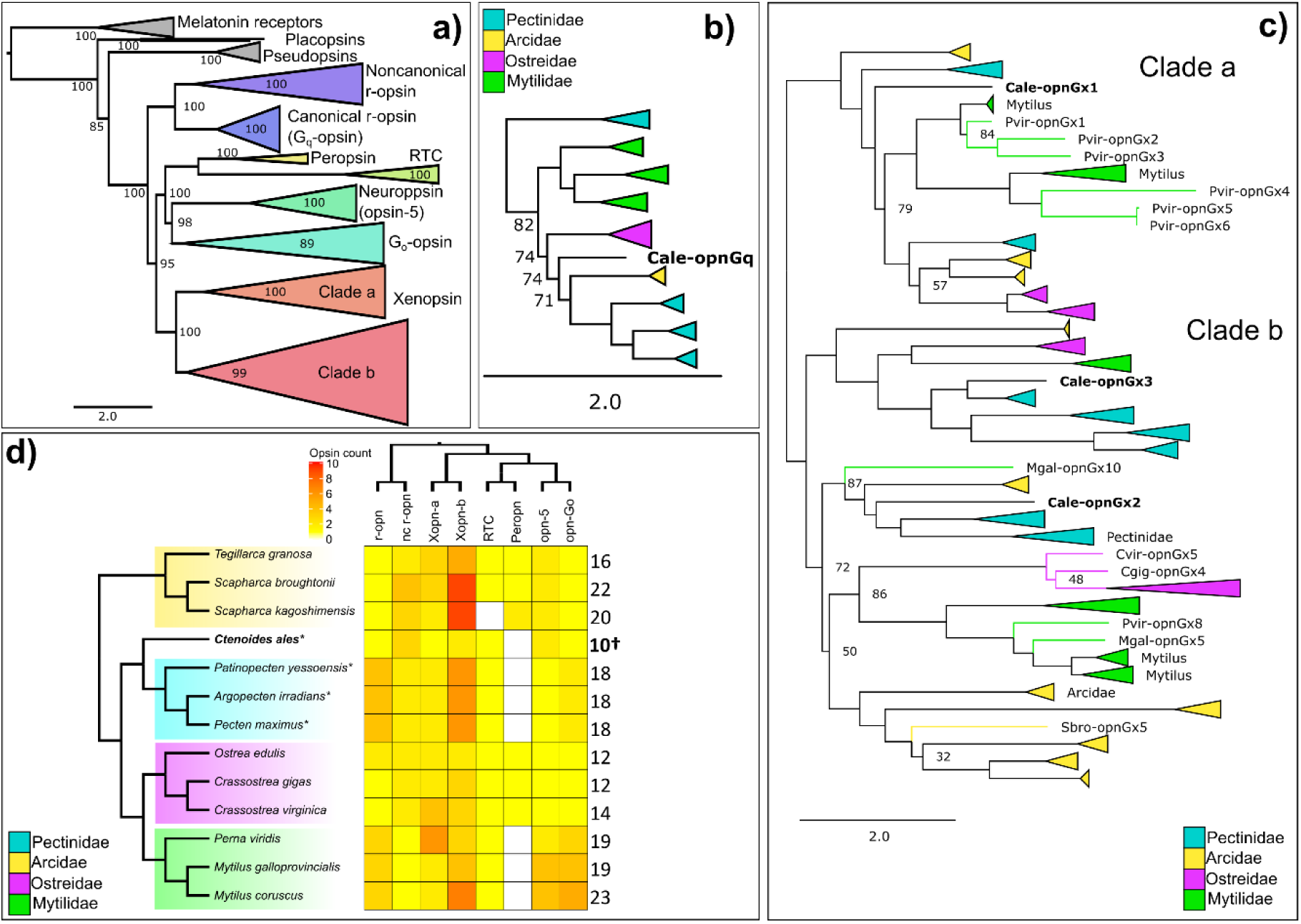
Summary of opsin content in *Ctenoides ales* and other pteriomorphian bivalves. **a)** ML phylogenetic tree of opsins from 13 pteriomorphian genomes, including *C. ales*. Ultrafast bootstrap (UFboot) support shown (Figure S4). Subtrees of **b)** “canonical” G_q_-coupled r-opsins (“opnGq”) and **c)** xenopsins (“opnGx”). In **b)** and **c)**, UFBoot support < 90 displayed; branches collapsed and color coded according to shared opsin duplicates within families; *C. ales* opsins noted in bold as “Cale-opn”. **d)** Heatmap reflects number of genes from each of 8 opsin groups (r-opn: rhabdomeric G_q_-coupled opsin; nc r-opn: noncanonical r-opsin; Xopn-a/b: xenopsin clades a and b; RTC: retinochrome; Peropn: peropsin; opn-5: neuropsin; opn-Go: G_o_-coupled opsin). Phylogenetic relationship among opsin groups based on Figure 3a. Total numbers of opsins per species reported right of heatmap. Shading around taxa according to family color coding in previous panels. *eyed species. †*C. ales* count includes the partial noncanonical r-opsin and G_o_-opsin.

The opsins in the *C. ales* genome belong to the major opsin groups present in mollusks: r-opsin (canonical and noncanonical), xenopsin, neuropsin, G_o_-opsin, and retinochrome (Ramirez et al. 2016) (Figure 3d).There does not appear to be a peropsin (Figure 3d) in the *C. ales* genome, which is also absent in Pectinidae and Mytilidae, but present in Ostreidae and Arcidae. The absence of perops in *C. ales* is likely a shared loss with Pectinidae. Compared to other species in Pteriomorphia, the limid, *C. ales* has a small opsin repertoire (Figure 3d). Unlike other families in Pteriomorphia, this representative limid lacks any lineage-specific opsin duplications (Figure 3d), even considering the partial noncanonical r-opsin and G_o_-opsin (Figure S4). The xenopsin group is particularly expansion prone in Pteriomorphia, as independent series of lineage-specific duplications are present in genomes from Pectinidae, Arcidae, Mytilidae, and Ostreidae. However, the three *C. ales* xenopsins are each phylogenetically located in separate clades of this opsin subfamily (Figure 3c), making lineage-specific paralogous duplication highly unlikely. Very little is known about xenopsins, as they were only recently recognized as a distinct group of opsins sister to the cnidarian “cnidopsins” (Ramirez et al. 2016). Xenopsins are only found in mollusks, other lophotrochozoans, and rotifers (Ramirez et al. 2016; Döring et al. 2020) and may be expressed in eyes along with r-opsins and c-opsins - depending on the species (Matsuo et al. 2019; Döring et al. 2020; Döring et al. 2020).

The “canonical” G_q_-coupled r-opsins are characteristically used in vision across the invertebrate clades of bilaterians. Previously, duplications of this opsin were identified in the bay scallop, *Argopecten irradians* (Porath-Krause et al. 2016), leading to the hypothesis that r-opsin expansion may be a common feature of eye evolution in bivalves. Here, we found only a single r-opsin in the *C. ales* genome versus the four in each of the scallop genomes (Figure 3b). This result indicates that scallop r-opsin duplication all occurred after the split from Limida. It also demonstrates that recruitment of additional r-opsins is not necessary for eye evolution in Pteriomophia. In fact, with eyeless mytilid species also having multiple r-opsin duplications (Figure 3b), the role of opsins in ocular vs. non-ocular processes requires particular attention in Pteriomorphia. The lack of opsin duplication in the *C. ales* genome contributes to the growing evidence for opsin evolution being unrelated to visual complexity in mollusks (De Vivo et al. 2023; McElroy et al. 2023).

Functional assays of *C. ales* opsins, including *in vitro* protein expression, as has been performed with scallop opsins (Smedley et al. 2022), as well as tissue-specific RNA-seq combined with *in situ* hybridization and/or immunohistochemistry will be valuable next steps in determining whether opsins from *C. ales* form photopigments and are expressed in eyes and other light-sensitive tissues.

### First mitogenome from Limida

Mollusks exhibit some of the most variable genomic architecture, molecular functions, and patterns of inheritance for mitochondria in metazoans (reviewed in Ghiselli et al. 2021). Pteriomorphian bivalves are known for dynamic mitochondrial (mt) genome evolution, with Arcidae having repeatedly evolved some of the largest bilaterian mt genomes (Kong et al. 2020), and Pectinidae with frequent gene order rearrangements (Malkócs et al. 2022) and species with exceptionally large mt genomes (e.g., La Roche et al. 1990). Currently, around 300 complete or nearly complete mitochondrial genomes are publicly available on NCBI GenBank, none of which are from Limida. In addition to being the first nuclear genome reported for a limid, we also present the first mt genome assembly for this order of bivalves, which should be a valuable resource for future phylogenetic analyses.

We assembled a 20,859 bp circular contig from the PacBio reads, representing a complete mitochondrial genome sequence. Using a combination of MITOS2, ARWEN, and ORF identification, we annotated 12 complete protein coding genes, the 12S and 16S rRNA genes, and 22 tRNA genes in the *C. ales* mitogenome (Figure 4). The only typical metazoan protein coding gene not annotated was atp8, which is commonly absent in bivalve mitogenomes (Serb and Lydeard 2003). However, recent analyses support the presence of atp8 sequences in Pectinidae (Malkócs et al. 2022) and Mytilidae (Zhao et al. 2022).

**Figure 4:**
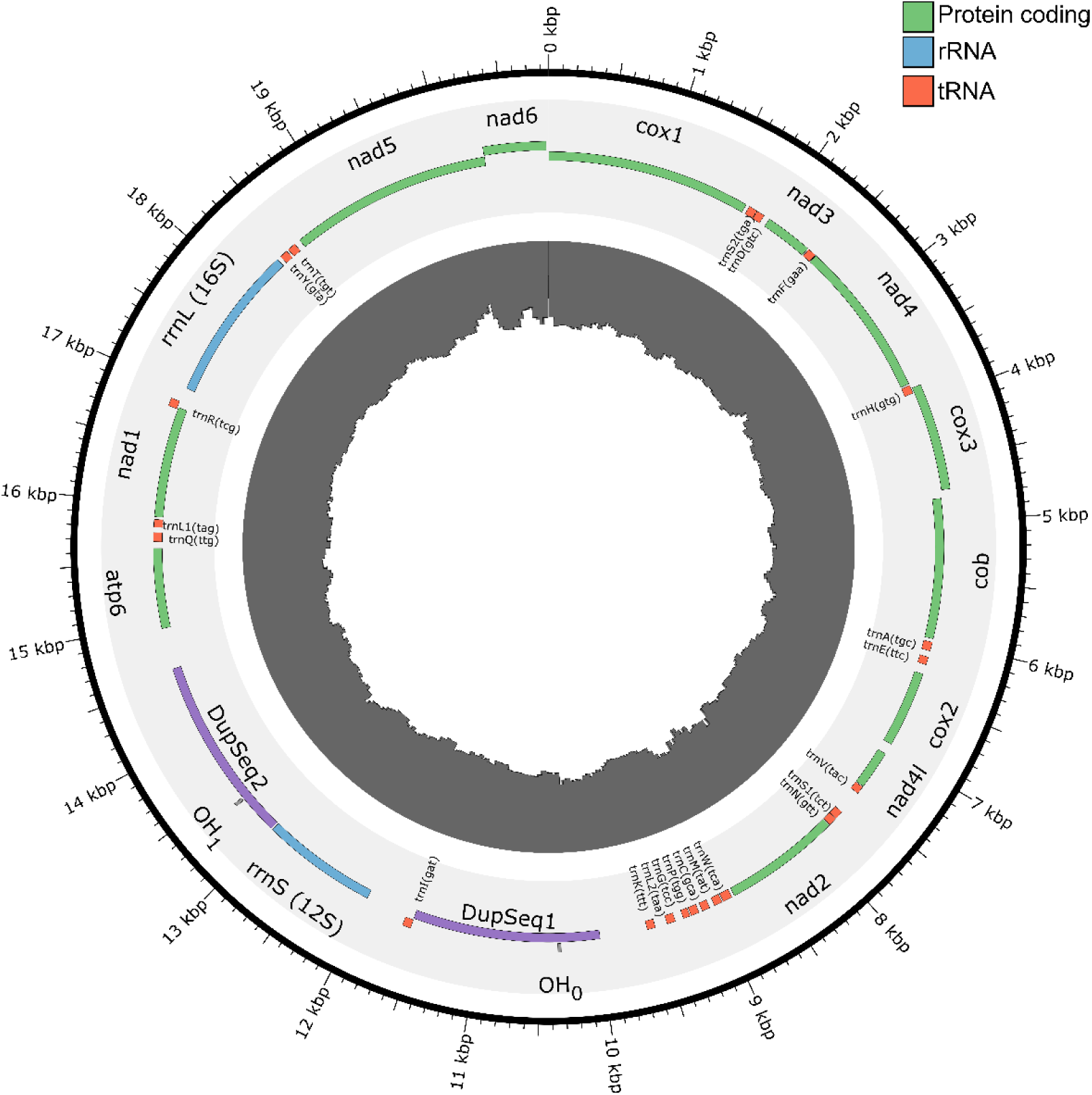
Annotation of *C. ales* mitochondrial genome. Protein-coding, rRNA, tRNA sequences color coding in legend, duplicated sequence with heavy-strand origin of replication (OH) in purple. Inner histogram of Illumina coverage (0-5282X), measured in 100bp windows step size of 50bp.

An interesting feature of this mitogenome is the apparent duplication of the heavy-strand origin of replication (“OH” annotation in MITOS, vs. “OL” for light-strand). We found a 1,589bp sequence duplicated on either side of the 12S rRNA gene that includes an OH annotation from MITOS. The two sequences are 99.8% identical, only differing at three positions. Many (907/4,230) PacBio reads span this repeated sequence (Figure S5) and we observed a 2X relative coverage of the repeat with Illumina reads mapped with one of the two sequences hard masked (i.e., all nucleotides reported as Ns) (Figure S6). These results support the accuracy of the mitogenome assembly in having a large duplicated sequence. This duplicated sequence containing OH likely represents the mitogenome “control region”, which regulates replication and transcription (Boore 1999). Duplication of control regions have occurred in a variety of taxa, e.g., birds (Schirtzinger et al. 2012), snakes (Jiang et al. 2007), and velvet worms (Braband et al. 2010), but little is known about the genetic architecture and/or evolutionary pressures underlying and maintaining duplicated control regions. The mitogenome of *C. ales* and limids more broadly may offer insight into the evolution of control regions and further supports bivalves as an emerging system for studying mitochondria (Ghiselli, Iannello, et al. 2021).

## CONCLUSION

In this study, we report a high-quality, chromosome-level assembly for *Ctenoides ales*, the first genome sequenced from the bivalve order, Limida. The genome of *C. ales* is noticeably larger than other pteriomorphian bivalves, largely due to a substantial number of transposable elements. We also find that this species has relatively few opsins, compared to other pteriomorphians, indicating that opsin diversification is not guaranteed to accompany the evolution of specialized adult eyes in bivalves. Additionally, we present the complete mitochondrial genome, another first for Limida.

## DATA AVAILABILITY

All raw read data have been uploaded to the NCBI SRA database, and the genome assembly has been submitted to NCBI GenBank. Raw reads and assembly are associated with the NCBI BioProject PRJNA1078364. The nuclear and mitochondrial genome assemblies with their annotations, the data underlying species and opsin phylogenies, and scripts used in this study will be available on Figshare doi:10.6084/m9.figshare.25290208.

## ACKNOWLEDGEMENTS

This project was supported by the National Science Foundation (DEB 1754331) to JMS and AJS and a Dovetail Tree of Life Grant Award to JMS. The research reported in this paper is partially supported by the HPC@ISU equipment at Iowa State University, some of which has been purchased through funding provided by NSF under MRI grants number 1726447 and MRI2018594. We are grateful to Justin Polson at Crowned Aquatics Pet Supply for providing care of the *Ctenoides* specimens. We thank Dr. Joel Sharbrough for helpful suggestions on analyzing the mitogenome.

